# A method for polyclonal antigen-specific T cell-targeted genome editing (TarGET) for adoptive cell therapy applications

**DOI:** 10.1101/2022.10.31.514542

**Authors:** Darya Palianina, Raphaël B. Di Roberto, Rocío Castellanos-Rueda, Fabrice Schlatter, Sai T. Reddy, Nina Khanna

## Abstract

Adoptive cell therapy of donor-derived, antigen-specific T cells expressing native T cell receptors (TCRs) is a powerful strategy to fight viral infections in immunocompromised patients. Determining the fate of T cells following patient infusion hinges on the ability to track them *in vivo*. While this is possible by genetic labeling of parent cells, the applicability of this approach has been limited by the non-specificity of the edited T cells.

Here, we devised a method for CRISPR-targeted genome integration of a barcoded gene into Epstein-Barr virus-antigen-stimulated T cells and demonstrated its use for exclusively identifying expanded virus-specific cell lineages. Our method facilitated the enrichment of antigen-specific T cells, which then mediated improved cytotoxicity against EBV-transformed target cells. Single-cell and deep sequencing for lineage tracing revealed the expansion profile of specific T cell clones and their corresponding gene expression signature. This method has the potential to enhance the traceability and the monitoring capabilities during immunotherapeutic T cell regimens.

## INTRODUCTION

Adoptive cell transfer of donor-derived antigen-specific T cells expressing native T cell receptors (TCRs) with defined specificities is an attractive immunotherapy strategy or clinical indications where polyclonality is beneficial. T cell therapies against cancer based on engineered TCRs or chimeric antigen receptors (CARs) typically only target a single antigen, reducing their applicable scope and making them vulnerable to relapses via antigen escape ^2^. In contrast, a polyclonal and polyspecific T cell population can target multiple antigens, potentially enhancing the overall effectiveness of an adoptive cell therapy ^3, 4^. The feasibility of this strategy has been demonstrated with virus-specific polyclonal T cells enriched from seropositive donors via stimulation with genetically-modified or Epstein-Barr virus (EBV)-transformed antigen-presenting cells (e.g., lymphoblastoid cell lines, or LCLs) ^5^ or rapidly expanded from peripheral blood mononuclear cells (PBMCs) using peptide pools as stimuli ^4, 6^, (manuscript in preparation). In this approach, single cell antigen specificity and phenotype characterization can be assessed prior to transfer through methods such as flow cytometry, ELISPOT and TCR RNA- or transcriptome-sequencing. These assessments become especially important during treatment. Beyond monitoring needs, the ability to identify the most therapeutically-relevant clones and phenotypes is of significant interest, particularly for long term efficacy. Recently, it was shown that CAR T cells can persist in patients as many as 10 years after infusion ^7^. While CAR T cells are readily identifiable, non-engineered therapeutic T cells are difficult to distinguish from naïve T cells. Genome-based lineage tracing of adoptively transferred lymphocytes has been proposed for facilitating follow-up studies ^8, 9^. For example, LCL-stimulated EBV-specific cells transduced with the *neo*-containing G1Na vector could be traced up to 9 years after adoptive transfer ^10, 11^. However, the use of retroviral vectors is associated with safety risks ^12^ due to the largely random nature of vector integration into the genome. Targeted gene editing by CRISPR/Cas9 is a superior approach and has been successfully used to knock out genes connected to exhaustion and checkpoint inhibition (e.g., PD-1) ^13^ or resist administered immunosuppressants (e.g., tacrolimus) ^14^. However, this approach has limitations, particularly for integrating a gene of interest, known as homology-directed repair (HDR). HDR is cell-cycle dependent and restricted to actively dividing cells ^15^. To date, CRISPR-based HDR approaches in T cells have relied on strong and nonspecific activation through anti-CD3 antibodies or coated beads. This approach is not compatible with a polyclonal T cell therapy where only target-specific cells are desired.

Here, we describe a novel approach for targeted CRISPR/Cas9-based genome editing and lineage tracing of virus-specific T cells. Notably, our approach combines autologous peptide presentation for T cell stimulation and editing, as well as the use of a barcoded GFP cassette library to enable the detailed characterization of clonal expansion. Using antigen-presenting cells and T cells directly from donor-derived PBMCs we generated a pool of uniquely barcoded EBV-specific T cells. By leveraging the cell cycle dependence of HDR, we used GFP integration as a marker of EBV specificity for enrichment by fluorescence-activated sorting (FACS). Sorted GFP-positive populations were devoid of unreactive cells as shown by single-cell RNA sequencing. This high purity resulted in an increased EBV-specificity and cytotoxicity against target cells (EBV-LCLs). Our method has a range of scientific and clinical applications: e.g., the possibility for sophisticated follow up after adoptive transfer on a single cell level, lineage tracing, the specific integration of therapy-enhancing genes such as a safety switch ^16^ or cytokines ^17^.

## RESULTS

### Library design and peptide-based T cell expansion

In order to fluorescently label and barcode reactive T cells in a single step, we designed an AAV vector encoding 1) inverted terminal repeats (ITRs); 2) homology arms for targeted insertion into the *CCR5* safe harbor locus ^18^; 3) the cytomegalovirus (CMV) constitutive promoter; 4) the GFP open reading frame (ORF) ; and 5) a 9-nucleotide barcode (Fig. 1a). Although the diversity of our library could theoretically encompass 262 144 unique barcodes, we restricted its size to 50000 colony-forming units. Sequencing of this cloned library identified 36030 unique barcodes (Fig. 1b) and no major bias (Fig. 1c). The repair template library was packaged into AAV6 capsid commercially and subsequently used for HDR following transfection.

**Fig. 1:**
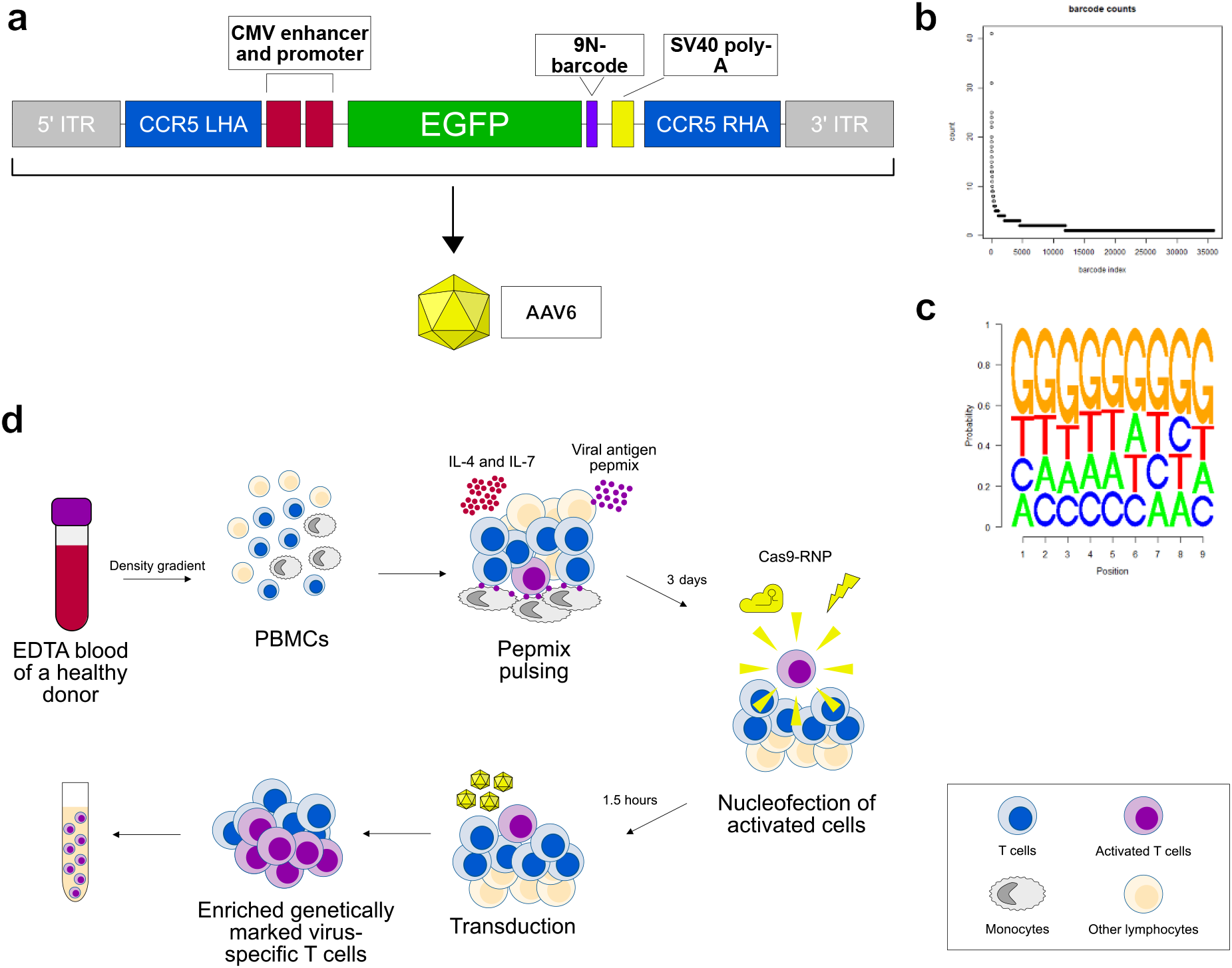
Library cloning and genome editing procedure. **a**, Library vector and repair template DNA design. Next generation sequencing (NGS) library analysis: **b**, sequencing visualization and **c**, sequence logo plots of the cloned library; **d**, Schematic of the cell culture and gene editing procedures. ITR – inverted terminal repeat, LHA and RHA – left and right homology arms, respectively; CCR5 – C-C chemokine receptor type 5, CMV – cytomegalovirus, SV40 polyA – simian vacuolating virus 40 polyadenylation signal, AAV6 – adeno-associated virus serotype 6, EDTA – ethylenediaminetetraacetic acid, PBMCs – peripheral blood mononuclear cells; RNP – ribonuclear protein.

To expand EBV-specific T cells from human PBMCs, we adapted an established protocol for rapid expansion of virus-specific cytotoxic T cells (CTLs) ^19^ and used the PepTivator EBV Consensus peptide pool as stimulus for display by native monocytes ^4^. This mixture covers 41 lytic and latent EBV antigens. It was previously shown that HDR is generally restricted to the S/G2 phases of the cell ^20^. Thus, proliferative activated T cells will preferentially undergo HDR following genome editing. We hypothesized that a population of pepmix-activated virus-specific T cells could be selected on the basis of successful HDR editing. To identify the optimal time-point for gene editing, we characterized the T cell proliferation and activation profiles of PBMCs from two healthy EBV seropositive donors with whole-cell staining and intracellular cytokine staining (ICC) every second day following EBV pepmix re-stimulation. No proliferation was observed among bulk T cells (Supplemental Fig. 1a) and EBV-responsive T cells (Supplemental Fig. 1b) by day 3, while daughter cells were present at day 5 and an abundant fraction of these were EBV-specific T cells. This lag between stimulation and expansion provided us with an opportune window for transfection. By transfecting prior to exponential cell expansion, we aimed to edit as many parent cells as possible. As such, we opted to transfect our barcoded library during this lag time, i.e., on day 3.

### Efficient transduction of peptide-stimulated T cells

In order to induce the genomic integration of a library in EBV-specific T cells, we devised the following strategy (Figure 1d). Following PBMC isolation from a healthy donor, cells were pulsed with EBV-pepmix or stimulated with anti-CD3/CD28 dynabeads in the presence of IL-4/IL-7. On day 3, cells were transfected with CRISPR/Cas9 ribonucleoprotein (RNP) and transduced with AAV6 particles carrying the barcoded GFP library. An RNP-only sample was included to serve as an HDR-negative control. On day 10, we analyzed cell type counts as well as GFP positivity. All samples including AAV-transduced were highly CD3+-enriched, confirming the efficiency of the pepmix and cytokines conditions for T cell enrichment (Supplemental Fig. 2). Cells transduced with the library showed GFP expression in both pepmix-stimulated and CD3/CD28 dynabeads-stimulated cells (Fig. 2a). Editing efficiency was donor-specific, ranging from 6.8% up to 25.0% for pepmix-stimulated cells and from 16.6% to 45.1% for bead-stimulated ones. For pepmix-stimulated product, we observed a higher proportion of GFP-positive T cells within the CD8+ population (median 12.5%) compared to those within the CD4+ one (median 5.7%), and we observed a similar trend for bead-activated T cells (medians 40.8% for CD8+ and 17.8% for CD4+). We also saw enrichment of CD8+ T cells in the AAV-transduced GFP-positive EBV-activated T cells compared to the bulk transduced ones (pepmix-stimulated but untransfected) product (p<0.05, 2-way ANOVA) (Fig. 2b).

**Fig. 2:**
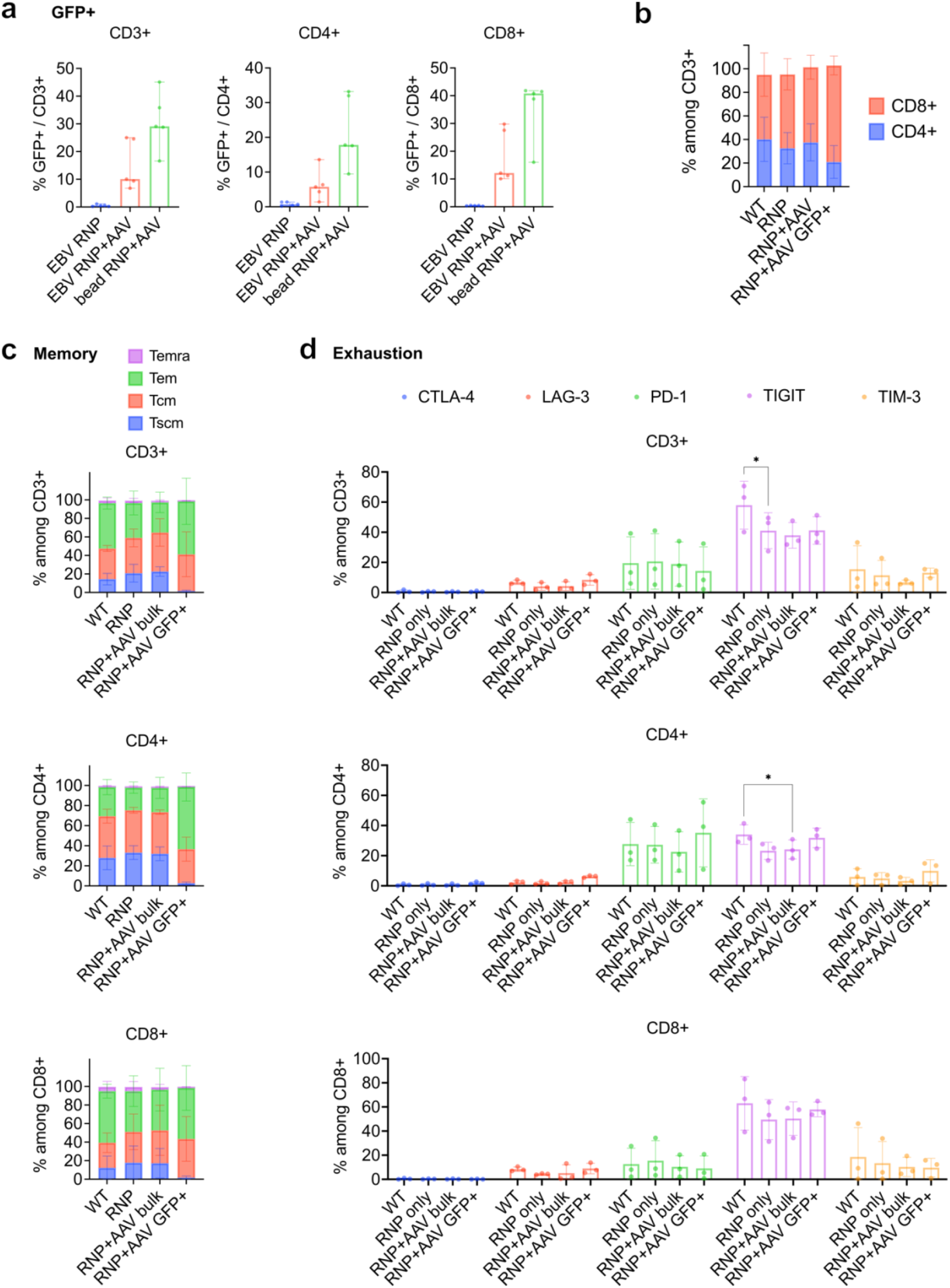
Transduction efficiencies and phenotype differences between expanded cells. **a**, Transduction efficiencies for pepmix-stimulated vs. anti-CD3/CD28 dynabeads-stimulated T cells for bulk CD3+, CD4+ and CD8+ cells, respectively; n=5, shown medians with range. **b**, CD4 vs. CD8 proportions within different populations of expanded pepmix-stimulated T cells; n=5, shown means with SD. **c**, memory phenotypes and **d**, exhaustion marker expression of expanded pepmix-stimulated T cells among WT, RNP-only transfected and transduced bulk CD3+, CD4+ and CD8+ cells; n=3. WT stands for wild type. Asterisk represent statistically significant differences (p<0.05, 2-way ANOVA). ANOVA – analysis of variance, EBV – Epstein-Barr virus, WT – wild type, RNP – ribonucleoprotein, AAV – adeno-associated virus; Temra – terminally differentiated, Tem – effector memory, Tcm – central memory, Tscm – stem cell memory T cells.

Next, we analyzed the memory phenotype of the generated EBV-CTLs (Fig. 2c). While untransfected samples showed an even mixture of stem cell memory (T_SCM_), central memory (T_CM_) and effector memory cells (T_EM_) with only a small minority of terminally differentiated (T_EMRA_) cells, we observed a depletion of T_SCM_ in GFP-positive T cells which comprised almost exclusively T_CM_ and T_EM_. This effect could be explained by low initial number of early-differentiated (T_SCM_ -like) EBV-CTLs in PBMCs due to EBV re-activation ^21^. Alternatively, early differentiated EBV-specific T cells might not be activated enough to enable HDR. Generally, CD4+ cells had a less differentiated phenotype compared to CD8+ in all conditions except among those GFP+-gated. Interestingly, among CD4+ GFP+ cells, there was a significantly higher proportion of T_EM_ compared to bulk transduced cells (p<0.05, 2way ANOVA).

We then assessed the expression of several exhaustion markers: CTLA-4, LAG-3, PD-1, TIGIT and TIM-3 (Fig. 2d). CTLA-4 was almost absent in all conditions, LAG-3 and TIM-3 were expressed at very moderate levels, slightly more among CD8+ cells compared to CD4+. PD-1 was overall also low but more present in CD4+ populations decreasing in AAV-transduced cells. On the contrary, TIGIT was expressed at high levels in CD8+ cells but less abundant in CD4+, decreasing further with both AAV transduction and RNP-only transfection. The decrease of TIGIT in transfected cells could be due to the death of exhausted cells following the transfection procedure.

Together, these results indicate that unique expansion and genome editing protocol efficiently integrated GFP in a population of activated T cells and did not markedly interfere with cell phenotype.

## HDR-based sorting enriches for EBV-CTLs and improves their anti-EBV response

In order to measure the EBV specificity and activation potential of the transduced bulk and GFP-positive cells, we re-stimulated expanded cells with pepmix and analyzed the expression of key cytotoxic T cell markers such as CD107a (LAMP-1), Granzyme B, IFNγ and TNFα using flow cytometry. While RNP-only transfected and bulk AAV-transduced cells did not show elevated cytotoxic marker expression compared to untransfected, within the GFP-positive T cell populations we saw elevated production of most markers (CD107a, IFNγ and TNFα) corresponding to at least a 2-fold increase in EBV specificity for CD8+ cells and 4-fold for CD4+ cells compared to wild type (Fig. 3a).

**Fig. 3:**
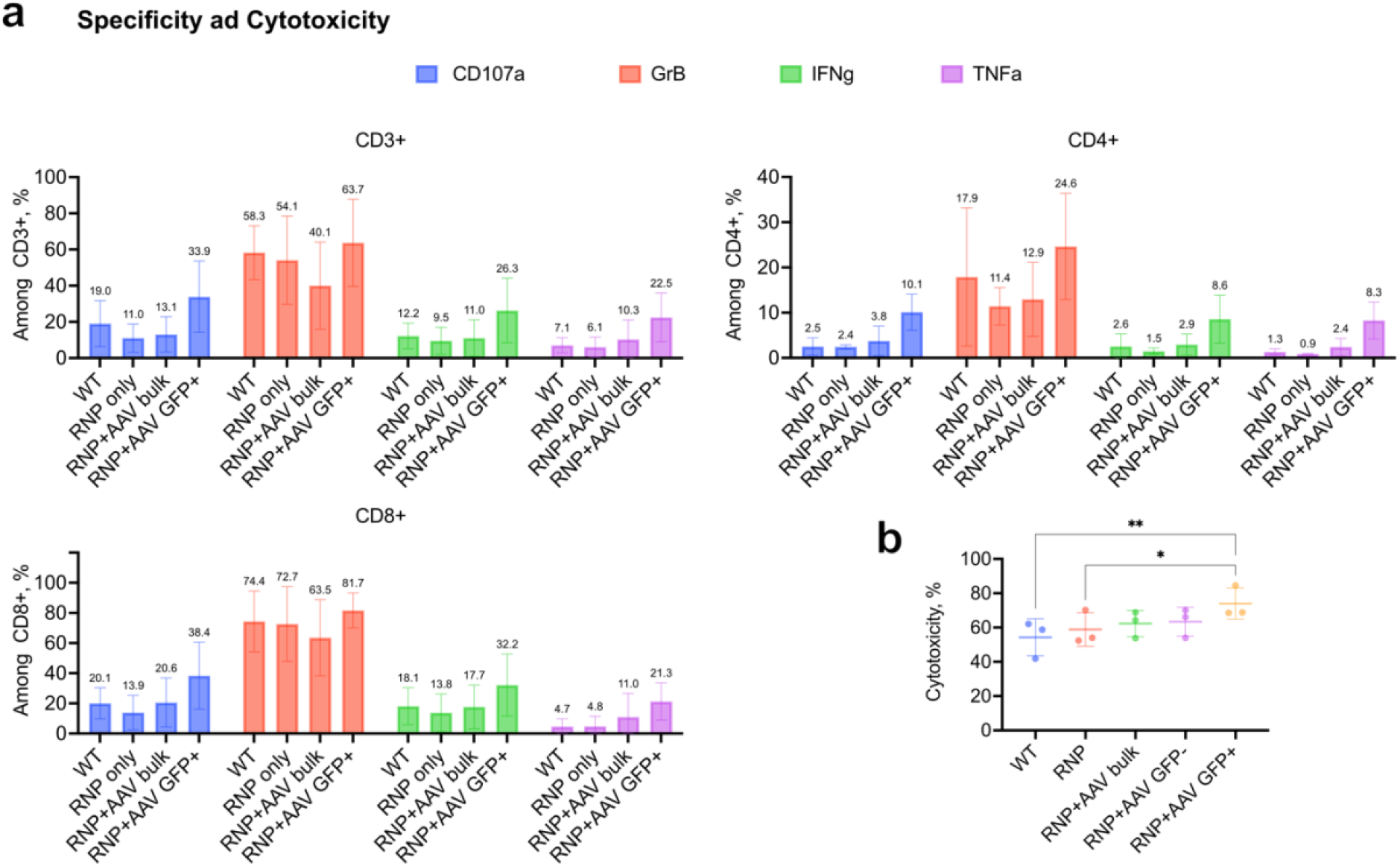
Specificity and functionality of pepmix-stimulated and expanded transduced T cells. **a**, Production of cytotoxicity markers and cytokines (CD107a, Granzyme B, IFNγ and TNFα) among bulk CD3+, CD4+ and CD8+ populations in response to EBV-pepmix-restimulation, n=3, means with standard deviation (SD). **b**, 6-hour cytotoxicity assay against autologous EBV-transformed LCLs (effector/target = 20:1), means of triplicates with SD for 3 donors, 2way ANOVA mixed effects analysis, **=0.0043, *=0.0405, α=0.05. ANOVA – analysis of variance, EBV – Epstein-Barr virus, WT – wild type, RNP – ribonucleoprotein, AAV – adeno-associated virus, GrB – granzyme B.

In order to assess target-specific functionality, we sorted GFP-positive and GFP-negative fractions of transduced EBV-CTLs and assessed their *in vitro* cytotoxicity against autologous EBV-transformed LCL and compared it to that of the other samples (Fig. 3b). Although we observed a slight increase of cytotoxicity in the RNP-only samples as well as the GFP-negative sorted fractions, this was less significant than that of the sorted GFP-positive cells highlighting the efficacy-enhancing potential of HDR-based selection.

These findings show that the designed HDR-based selection of EBV-CTLs leads to an increased antigen specificity and target-specific toxicity.

### GFP barcoding and selection provide expansion and enrichment statistics, respectively

Ten days following peptide pulsing, we sequenced 38 908 cells across two donors and the two sorting conditions (GFP-positive and GFP-negative) from which 27 283 had a properly annotated TCR (Fig. 4a). Single cell sequencing provided us with three layers of data for both edited and unedited T cells: 1) TCR clonal identity; 2) GFP barcode clonal identity; and 3) Gene expression (transcriptome) profiles. Among all cells, 295 unique TCR clonotypes appeared at least three times. One highly represented clonotype, representing 45% and 65% of the GFP-positive and -negative datasets for Donor 2, respectively, was omitted for TCR identity analysis as a likely indiscriminately-expanding clone. Of the remaining clonotypes, none were shared between donors, while V and J gene usage diversity was also distinct (Fig. 4b).

**Fig. 4:**
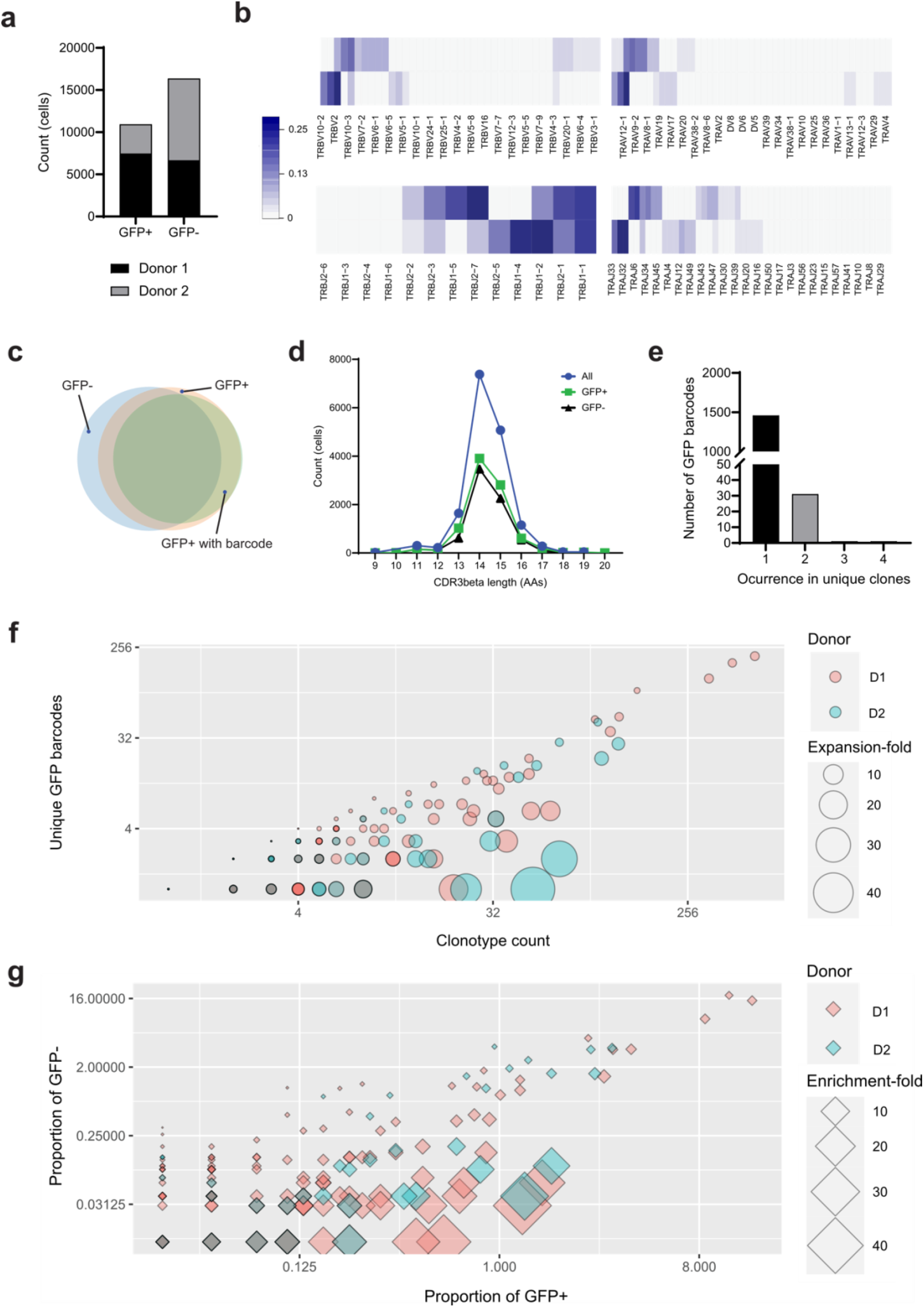
Lineage tracing and enrichment analysis of single-cell sequencing data reveals highly-expanded and highly-enriched TCR clones. **a**, Sequenced cell counts and their sample and donor origin. Each donor was used in a single expansion, genome editing, sorting and sequencing workflow. Cells were sorted for the presence (GFP+) or absence (GFP-) of GFP fluorescence. **b**, Heatmaps comparing the sequenced alpha and beta TCR chains and their V- and J-gene usage for each donor. Donors showed markedly different gene usage profiles. **c**, Venn diagram of the membership of TCR clonotypes across GFP-positive and GFP-negative samples. The high overlap between the samples enabled the downstream calculation of enrichment statistics. **d**, Distributions of the CDR3 beta lengths for GFP+ and GFP-samples. The distributions do not differentiate between samples. **e**, Frequencies of identical GFP barcodes found in more than one T cell clone, based on TCR identity. The vast majority of GFP barcodes were associated with only a single clone, confirming that the barcode library was sufficient. **f, and g**, Scatter plots of the fold-expansion and fold-enrichment respectively for individual clonotypes. Fold-expansion was calculated from the ratio of GFP barcodes to clonotype count. Fold-enrichment was calculated from the ratio of proportions within libraries, from GFP- to GFP+. Grey data points show perfect overlap across donors. Both statistics enable clonotype comparisons.

We next compared the GFP-positive and -negative sorted samples. Clonotype overlap between samples was high (Fig. 4c), while CDR3 length distribution were a close match (Fig. 4d, Supplemental Fig. 3a). To better distinguish between clonotypes, we investigated post-GFP integration fold-expansion (cell proliferation after day 3) and fold-enrichment (selection efficiency). For expansion, we performed a lineage tracing analysis through deep sequencing of the GFP gene to specifically link cell and GFP barcodes. We obtained GFP barcodes for 44% of the sequenced GFP-positive cells, representing 1491 unique barcodes. Only a minor fraction of these (2%) were associated with more than one TCR clonotype (Fig. 4e) and were excluded from subsequent analyses. Using the ratio of GFP barcodes to cell barcode, we calculated the mean fold-expansion of 209 individual T cell clonotypes (Fig. 4f). While the highest expansion was 49-fold (with one GFP barcode), the middle 50% of clones ranged between 1- and 3-fold. Interestingly clonotypes with the highest post-GFP integration fold expansion did not correlate with the clonotypes that had overall the highest number of cells, revealing interesting clonotype expansion dynamics.

In addition to fold-expansion, we calculated fold-enrichment for the 170 clonotypes that were assigned to a GFP-barcode and had cells in both GFP-negative and -positive samples, based on their enrichment across samples (Fig. 4g). Selection on this basis resulted in enrichment as high as 43-fold, or in depletion as high as 12-fold. Fold-expansion and fold-enrichment showed a significant correlation (P < 0.0001) though perhaps driven by a handful of clonotypes (Supplemental Fig. 3b).

We submitted the beta chains’ V gene, J gene and CDR3 for TCRs with barcoded GFP to TCRex, a tool designed for querying TCR identity in public databases (TCRex (biodatamining.be)). Fourteen clones within eleven clonotypes were classified as EBV-specific, three of which showed enrichment and expansion both above one (Table 1). Of these, two are perfect matches by CDR3β to dominant clones highlighted in previous work ^22-24^, while clonotype 110 is a close match (Levenshtein distance of 3). EPLPQGQLTAY and GLCTLVAML correspond to peptides from BMLF1 and BZLF1 lytic EBV proteins while HPVGEADYFEY (EBNA1) and IVTDFSVIK (EBNA4) correspond to the latent ones. Overall, we have high confidence that we identified multiple EBV-specific TCRs for which we have lineage tracing data.

**Table 1:**
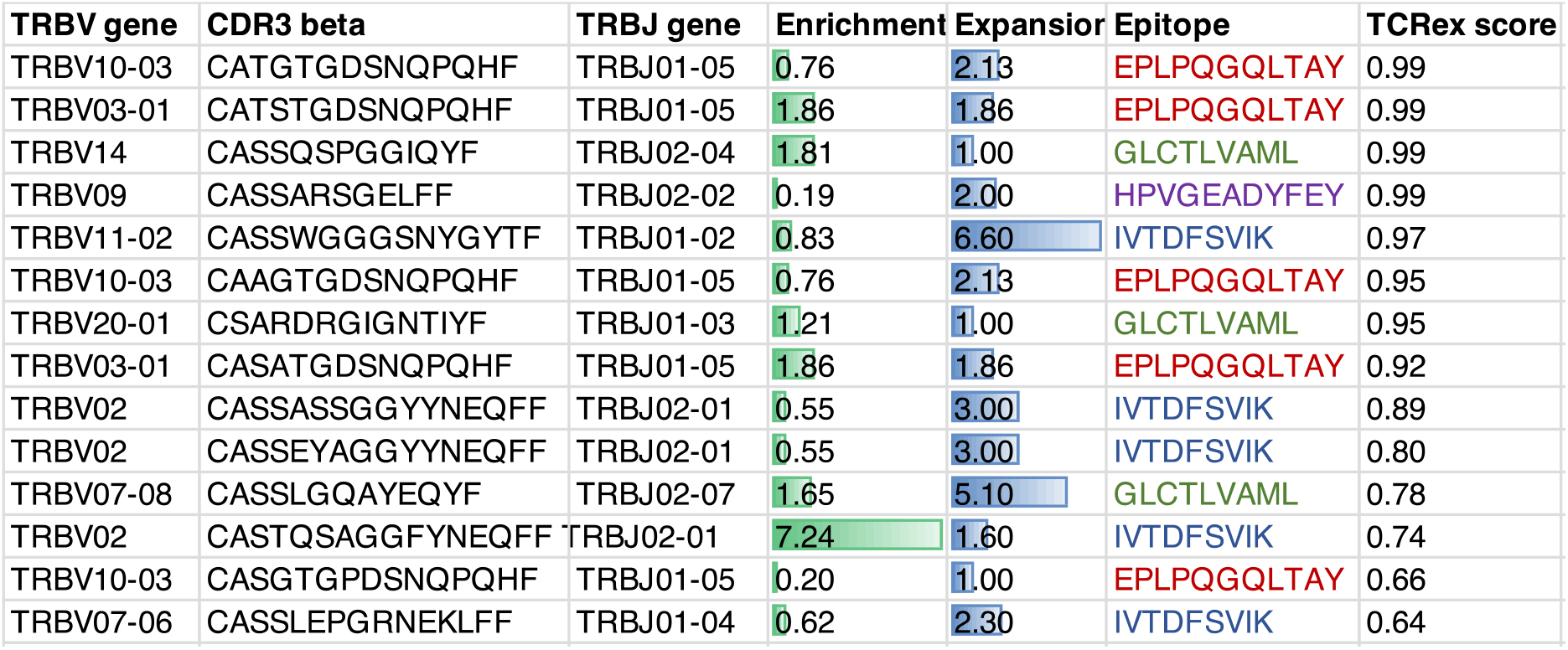
EBV-specific TCR clones within GFP-positive and barcoded dataset as predicted by TCRex.

### Single-cell transcriptome sequencing confirms the enrichment of reactive T cell phenotypes in GFP-positive sorted cells

Using single-cell transcriptomics, we explored the phenotypic landscape of EBV-CTLs. Unsupervised cell clustering divided all cells into 13 main clusters (Fig. 5a-b and Supplemental Fig. 4a-c). With few exceptions, CD4/CD8 identity, cell cycle phase, cytotoxicity and memory markers were the main drivers of cluster separation. CD8 clusters 0,1,2,3,5,7 and 8 describe a homogeneous population of activated cytotoxic CD8 cells enriched in the expression of NKG7, GZMK, GZMA, GZMH, PRF1, HLA-DRA, and EOMES. Clustering resolves cycling cells (clusters 1 and 2), non-proliferative CD27/CCL4/CCL5-high and GZMB/LAG3-high cells (clusters 0 and 3 respectively), glycolytic cells (cluster 5) and apoptotic cells (cluster 8). Cluster 4 is a CD4-enriched cluster of moderately proliferative cells presenting an activated phenotype and retaining the expression of memory markers such as TCF7, LEF1 and CD7. T reg CD4 cells are found in cluster 9 enriched in FOXP3/ ILR2A and lastly, cluster 6 describes a CD4/CD8 population of resting cells enriched in memory and resting cell markers such as IL7R, CCR7 and TCF7 which present a phenotype of unreactive T cells. Clusters 10, 11 and 12 show small populations of NK and B cells remaining from the initial whole-PBMC population.

**Fig. 5:**
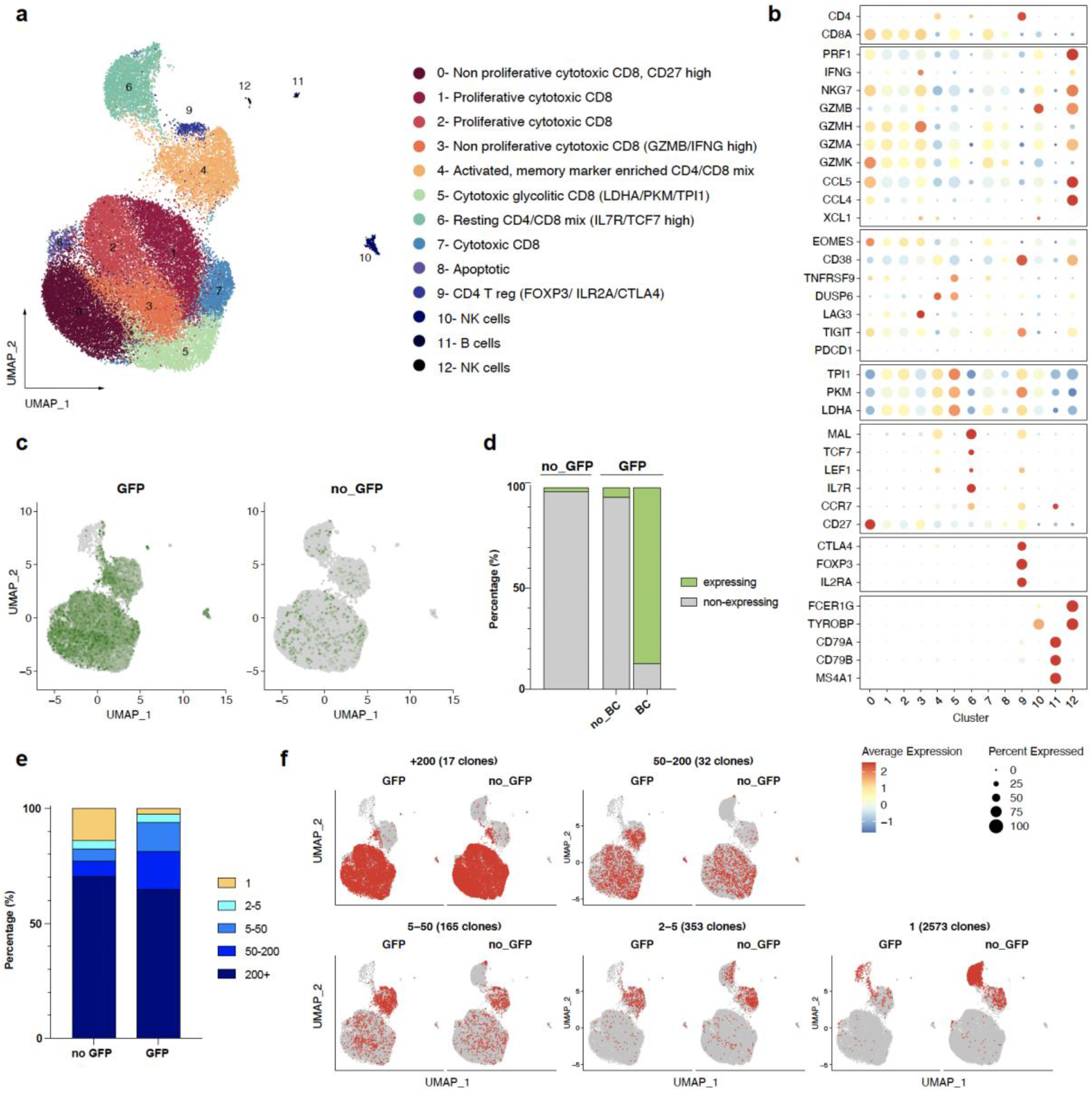
Single cell transcriptomics and TCR sequencing reveals a broader enrichment of EBV reactive T cell phenotypes in GFP sorted cell samples. **a**, UMAP embedding and unsupervised cell clustering of 38908 EBV pulsed T cells. **b**, Dot plot showing the expression of a selection of T cell marker genes across clusters found in A. **c**, Feature plots showing the distribution of GFP expression across cells from GFP positive and GFP negative sorted samples. **d**, Enrichment of GFP positive and GFP negative sorted sample groups in GFP expressing cells. GFP positive sample group is further divided into cells that did or did not have a correctly annotated GFP-barcode. **e**, Enrichment of GFP positive and GFP negative sorted sample groups in 5 different TCR clonotype expansion bins. **f**, Projection of cells from 5 different TCR clonotype expansion bins on to the transcriptomic UMAP embedding. Cells from GFP positive and GFP negative sorted sample groups are shown in separate UMAP plots. BC – barcode.

The detection of GFP transcripts at single cell level was used to confirm the correct CRISPR-Cas9 genome integration of the GFP-Barcode transgene (Fig. 5c-d). About 45% of GFP-positive sorted cells showed detectable GFP transcript, compared to just 2% for the GFP-negative sample. Moreover, 87% of the GFP-positive sorted cells that were assigned to a correctly annotated GFP barcode by deep sequencing also expressed GFP transcripts, further demonstrating the accuracy of our GFP barcode readout.

When comparing the enrichment of GFP-positive and -negative cells across clusters, GFP-positive cells were strongly depleted in cluster 6 (unreactive T cells) and enriched in activated T cell clusters 1 through 5 (Fig. 5c and Supplemental Fig. 5a-b). In addition, we saw different patterns of enrichment when looking at the expansion of TCR clonotypes within these two sample groups. Cells sorted for absence of GFP were enriched for non-expanded (only one cell per clone) and highly expanded (more than 200 cells per clone) clonotypes as opposed to GFP sorted cells, which were enriched in highly expanded and also moderately expanded clonotypes (5-200 cells per clone; Fig. 5e-f). Cluster enrichment across TCR expansion bins and the top expanded clonotypes showed that overlooking moderately expanded clonotypes restricted the diversity of T cell phenotypes (Supplemental Fig. 6a-b). On the other hand, our results showed that this could be avoided by using our targeted GFP-positive selection; while most TCR expanded clonotypes clustered around the same phenotypes, the expansion of our GFP-barcode was distributed more homogeneously across activated T cell phenotypes (Supplemental Fig. 7).

These results illustrate how our method can be more effective in identifying highly but also moderately expanded reactive T cells across any activated phenotype for both the CD4 and CD8 compartments.

## DISCUSSION

Adoptive T cell therapy is a highly versatile treatment option due to the involvement of T cell immunity in a variety of indications such as autoimmunity ^25^, blood ^26^ and solid^27^ cancers, infectious diseases ^28, 29^ and diabetes ^30^, to name but a few. While CAR-T cells are becoming a standard-of-care treatment for some hematological malignancies, patients with other challenging indications would benefit from alternative options with enhanced efficacy and persistence or with a broader targeting spectrum such as those afforded by the use of isolated antigen-specific T cells with native TCR, as shown for virus-associated malignancies ^18^.

Here, we present an efficient and polyvalent method of targeted gene delivery into antigen-specific T cells using a CRISPR protocol adapted to the use of peptide antigens as HDR-enabling stimuli in contrast to the commonly used nonspecific anti-CD3/CD28 stimulation of T cells. Although we focused on EBV-CTLs as a proof of concept, we note that this method does not depend on the specifics of this model, and can therefore also be applied to other antigen targets such as anti-tumor WT1-reactive T cell enrichment for anti-leukemic activity ^31^.

The use of genome editing for cell engineering offers notable advantages, in particular the precision of DNA construct integration. This ensures minimal disruption of cell function, as evidenced by our post-transfection phenotypic analysis, as well as long-term lineage tracing. While a typical weakness of CRISPR-induced HDR lies in its efficiency, we were able to achieve rates of integration suitable for a substantial DNA barcode library. Coupled with the permanence of genome editing, DNA barcodes may soon become standard procedure in cell therapies ^32^, making next-generation sequencers a likely soon-to-be essential tool in the clinic. In order to assess the quality of the barcoded and selected EBV-CTLs, we combined our methodology with scRNA-seq, another tool that is revolutionizing cell engineering and immunotherapies ^33^. Single-cell barcode sequencing, coupled with TCR clonotype information, provided an unprecedented level of detail on clonal expansion. In addition, scRNA-seq can link lineages to specific T cell phenotypes. The heterogeneity of stimulated T cell populations is essential to the development of effective immunity, and our genome editing protocol does not interfere with phenotype diversity. For instance, beyond the cytotoxic potential of CD8 T cells, it has been clearly shown that CD4+ T cells are crucial for sustaining anti-viral memory and effector functions ^34, 35^. We observed that antigen-specific T cell stimulation combined with genome editing-based selection enabled the enrichment of EBV-CTLs with both CD4 and CD8 populations showing increased production of CD107a and cytotoxic molecules such as Granzyme B, IFNγ and TNFα among GFP-positive cells. Memory composition is another critical parameter of an effective therapeutic T cell product ^36^. Early differentiated memory phenotypes such as stem cell memory and central memory are superior in the sustaining long-term anti-tumor responses ^37, 38^. Generally, we had high proportions of central memory population among the GFP-positive cells and a good enrichment of memory markers in the transcriptomics of the GFP-sorts. Interestingly, we observed a decrease of naïve-like/stem cell memory like CD62L+CD45RA+ population in contrast to bulk transduced or wild type cells which could be explained either by a slower activation of early-differentiated cells compared to central and effector memory cells and as a result lower level of HDR, or by initially low amount of EBV-specific T cells among early-differentiated cells due to a high frequency of EBV (CTL-cell-controlled) reactivation in humans ^21^. Functionally, we noted that GFP-positive sorted T cells exhibited enhanced antigen specificity and improved cytotoxicity against autologous EBV-transformed LCLs.

Our work constitutes the first instance of the precise introduction of a genetic marker targeting selected donor-derived antigen-specific T cells. The method and these data combined should help establish the next generation of cell therapies combining *in-vitro* and *in-vivo* lineage tracing and the functional enrichment of antigen-specific T cells.

## METHODS

### Plasmid library construction

The barcoded GFP library was encoded in a plasmid constructed in two steps. First, the pCMV-GFP and homology arms were designed *in silico* and synthesized externally (Twist Bioscience). Second, the GFP was barcoded using oligonucleotide F1(RB203)* (Supplemental Table 1) with 9 degenerate “N” nucleotides and flanking regions homologous to the end of the GFP open reading frame and the start of the polyA signal. The oligonucleotide was used with primer R1(RB202)* in a NEBuilder assembly reaction (NEB). The resulting plasmid was transformed in electro-competent *E. coli* DH5α cells which were then grown in Luria-Bertani broth with 50 μg/ml ampicillin. An aliquot was plated to assess the transformation efficiency.

### Peripheral blood mononuclear cell (PBMC) extraction and cell culture

EDTA blood collected from adult healthy donors was used for peripheral blood mononuclear cell (PBMC) extraction. The study was approved by the Ethical Committee of Northwestern and Central Switzerland (PB_ 2018-00081), and written informed consents were obtained. PBMCs were isolated as previously published^39^. All cells were cultured at 37°C, 5% CO2.

T cells were cultured in cytotoxic T cell line medium (CTLm) composed of RPMI (Gibco), 5% human serum and 10,000 U/mL Penicillin-Streptomycin (ThermoFisher). PBMCs were stimulated with either anti-CD3/CD28 Dynabeads (ThermoFisher) according to manufacturer’s instructions or with EBV pepmix (PepTivator EBV Consensus peptide pool (Miltenyi Biotec)), at a final concentration of 60 pmol/peptide/mL in CTLm supplemented with 400 U/mL IL-4 and 10 ng/mL IL-7 (R&D Systems) for three days. After that, cells were washed, transfected and cultured n CTLm with cytokines or cultured without transfection.

EBV-transformed LCLs were generated using the B95.8 EBV strain as previously published 40.

### Cell proliferation assay

1.5×10^7^ PBMCs were stained with CellTrace™ Violet (CTV) Cell Proliferation Kit according to manufacturer’s protocol, stimulated with the EBV pepmix and cultured in 6-well GRex plates (Wilson Wolf) and cultured for 9 days. Every second day starting day 3 of culture cells were gently resuspended and a fraction of cells was taken for immunocytochemistry and cell proliferation tracing by flow cytometry.

### Genome editing of EBV-specific T cells

PBMCs were genome-edited using a combination of CRISPR/Cas9 ribonucleoprotein (RNP) and adeno-associated viral particles (AAV) after three days of culture with or without stimulation. The RNP was assembled by first duplexing the CRISPR RNA (crRNA, sequence TGACATCAATTATTATACAT CGG ^41^) and trans-activating CRISPR RNA (trcrRNA) (IDT) through co-incubation at 95°C for 5 minutes and cooling to room temperature. The duplexed RNA molecules were then complexed with 25 μg (153 pmol) of Cas9 protein (IDT) at room temperature for 20 minutes. The AAV particles were produced externally (Vigene Biosciences) by packaging the repair template DNA encoding the pCMV-barcoded GFP construct in a AAV6 capsid. From the PBMC cultures, cells in suspension were gently extracted without a detaching agent. The culture wells, which retained adherent monocytes, were gently washed and topped with serum-free CTL and set aside during the transfection procedure. Suspension cells were centrifuged to remove the culture medium and resuspended in 100 μL P3 nucleofection buffer (Lonza), to which 6.5 μL of RNP were mixed in. Cells were transferred to nucleocuvettes and shocked using a 4D-Nucleofector (Lonza) with protocol EO-115. Cells were then gently diluted in 600 μL of warm serum-free CTL medium. After 30 minutes, the transfected cells were placed in their original well after emptying them again without detaching monocytes. After two hours of incubation, 20 μL of AAV particles at 2.25×10^13^ particles/mL (for a target MOI of 2×10^5^ particles/cell) were added to the cultures. After 24 hours, the cultures were diluted 1:1 with human serum-supplemented CTL medium.

### Fluorescence activated cell sorting (FACS) of GFP+ cells

Expanded EBV-stimulated and transduced T cells were sorted based on GFP fluorescence after 10 days of culture. Cells in suspension were gently extracted without a detaching agent and centrifuged to remove the culture medium. Cells were then washed in DPBS (Gibco), sorted using SH800 cell sorter (Sony Biotechnology) into CTL medium. For specificity and cytotoxicity analysis, cells were recovered for three days in CTLm supplemented with IL-4 and IL-7.

### Staining for flow cytometry analysis of surface markers

Cells were stained with Zombie Aqua viability dye (Biolegend) in PBS, washed in FACS buffer and stained with the cocktail of monoclonal antibodies for CD3-BUV395 (clone UCHT1), CD4-BUV496 (SK3), CD8-BUV805 (SK1), TIM-3-BV480 (7D3), PD1-BB700 (EH12.2H7) (all BD Biosciences); CD45RA-APC (MEM-56, Thermo Fisher Scientific); CD45RO-Alexa Fluor 700 (UCHL1), CD62L-BV650 (SK11), CD27-BV421 (M-T271), CTLA-4-BV785 (L3D10), LAG-3-BV711 (11C3C65, Biolegend) and TIGIT-BV605 (A15153G, Biolegend).

### Intracellular cytokine staining (ICC)

Cells in a pure CTLm as a negative control and cells stimulated with EBV pepmix were seeded into a U-bottom 96-well plate containing pure CTLm as a negative control, or CTLm with 500x-diluted pepmix, respectively. Cross-linked costimulatory anti-CD28/CD49d monoclonal antibodies (BD Biosciences), 1 μg/ml each, and anti-CD107a-BV510 (H4A3, Biolegend) were added, and cells were incubated at 37°C, 5% CO2 for 1 hour. Next, cell transport was blocked by 10 μg/ml Brefeldin A (Sigma). 5-hour incubation was followed by intracellular staining for flow cytometry analysis.

Cells were stained for viability with Zombie UV dye (Biolegend) according to manufacturer’s instructions. Next, cells were washed with FACS buffer (2% sterile filtered FBS and 0.1% NaN3 in PBS), stained with surface monoclonal antibodies (all from BD Biosciences) for CD3-BUV395 (UCHT1), CD4-BUV496 (SK3), CD8-BUV805 (SK1) in FACS buffer, washed, fixed with fixation buffer (Biolegend), washed with permeabilization buffer (Biolegend) and stained for 30 min with the cytokine antibodies for (all Biolegend): IFNγ-APC/Cy7 (B27), TNFα-PE/Cy7 (MAb11) and Granzyme B-PE/Cy5 (QA16A02).

### Cytotoxicity assay

T cells were incubated with autologous LCLs (Effector:Target = 20:1) for 5 h 30 min, stained for apoptosis with CellEvent Caspase-3/7 (Thermo Fisher), incubated for additional 30 min, washed in PBS, stained for dead cells with Zombie Aqua (Biolegend), then stained for CD3+ and CD19+ in FACS buffer and analyzed by flow cytometry. LCLs incubated without T cells were used as a control. The analysis was performed as previously published ^4^. The formula used to define cytotoxicity was: % specific lysis = 100 – ([Vtest/Vcontrol]*100), where V is percentage of viable cells (double-negative for ZombieAqua and CellEvent).

### Flow cytometry analysis

Samples were acquired on Cytek Aurora using SpectroFlo software. Data were processed using FlowJo.

### Statistical analysis of flow cytometry data

Data were analyzed in Prism (GraphPad) using ANOVA or 2way ANOVA via statistical methods whichever were applicable.

### Genomic PCR

Genomic DNA from 10^4^ to 10^5^ T cells was was extracted using QuickExtract buffer (Lucigen). The resulting product was used as a DNA template for a first PCR amplification reaction using primers F2(RB198)* and R2(RB199)* primers (Supplemental Table 1). The 3 kbp product was extracted by gel agarose electrophoresis and used as template for a second PCR amplification using primers F3(RB214)* and R3(RB215)*. The final amplimers were purified and sequenced externally by Illumina paired-end sequencing (GENEWIZ).

### Single-cell sequencing

Single-cell sequencing was done according to the 10X Genomics pipeline and the manufacturer’ instructions as previously described ^42^. Briefly, for each donor, 20000 GFP-expressing cells and 20000 GFP-negative cells were sorted as described above and processed for single-cell sequencing using a Chromium Next GEM Single Cell 5’ Library & Gel Bead Kit v1.1, a Chromium Next GEM Chip G Single Cell Kit and a Chromium Controller. The gene expression (GEX) and the fragmented TCR VDJ targeted enrichment libraries were prepared using a Chromium Single Cell 5’ Library Construction Kit and a Chromium Single Cell V(D)J Enrichment Kit, Human T Cell. For GFP targeted enrichment, the primers F4(RB200)* and R4(RB222)* were used in a PCR amplification reaction. The resulting product was used as template for a second PCR amplification using an indexing primer and primer R5(RB201)*. All libraries were indexed using primers from a Chromium i7 Multiplex Kit and sequenced by the Genomics Facility Basel using an Illumina Novaseq and a single lane of a S4 flow cell.

### Analysis of scRNA-seq GEX data

The raw scRNA-seq data was aligned to the human genome (GRCh38) using Cell Ranger (10x Genomics, version 6.0.0). In the first place a custom reference human genome, incorporating the GFP gene, was created using the *mkref* function, then the *count* function was used to obtain the raw gene expression matrix. Downstream analysis was carried out using the Seurat R package (version 4.0.1). As quality control, cells presenting low and high number of detected UMIs (200 < nFeature_RNA < 7,000) and high percentage of mitochondrial genes (Percentage_MT < 20% of total reads) were removed. In addition, TCR genes were removed to avoid clonotype from guiding the subsequent clustering.

After QC a total of 38908 cells were used for downstream transcriptomic analysis. All samples were merged, normalized and scaled using 2000 variable features (GFP gene was removed to avoid its influence in downstream clustering analysis) while also regressing out cell cycle scores. Dimensionality reduction was done using the *RunPCA* function and batch effect was removed by performing harmony integration. Finally unsupervised cell clustering and differential gene expression was used to find marker genes used for cluster annotation. Results were then visualized using UMAP dimensionality reduction and ggplot2 R package.

### Paired TCR repertoire analysis

Raw TCR scRNA-seq data was aligned to the VDJ-GRCh38-alts-ensembl (5.0.0) using Cell Ranger (10x Genomics, version 3.1.0). As quality control, only cells retaining a productive alpha and a productive beta chain were used. Downstream analysis was done using the R programming language and common packages (code available upon request). Cluster definition was performed as previously described by ^42^ and the comparisons of V and J gene usage was done using the package bcRep ^43^.

### Analysis of scRNA-seq GFP barcode data

The GFP barcodes were linked to single cells through the 10X Genomics barcode. Where two GFP barcodes were identified (likely bi-allelic integration), they were concatenated and treated as one. For EBV-specificity predictions using TCRex, V- and J-gene information, along with CDR3 beta sequences, were queried against all available EBV epitopes. The output was then re-linked to clonotype identity.

## Supporting information

TarGET_Supplemental

## ACKNOWLEDGEMENTS

We thank the Genomics Facility Basel and FACS Core facility of the Department of Biomedicine, University Hospital of Basel, for excellent support and assistance throughout this study. Acknowledgements go to the blood donors, and to Dr. Glenn Bantug for providing the EBV strain. This work was supported by the Swiss National Foundation Grant 32003B_204944 (to N.K.), NCCR Antiresist Grant No. 180541, Switzerland (to N.K.), Bangerter–Rhyner Stiftung (to N.K.), ETH Zurich Post-doctoral Fellowship, Switzerland (to R.B.D.R.), Helmut Horten Stiftung, Switzerland (to S.T.R.) and NCCR Molecular Systems Engineering, Switzerland (to S.T.R.).

## AUTHOR CONTRIBUTIONS

D.P., R.B.D.R, S.T.R. and N.K. designed the study; D.P., R.B.D.R. and R.C.R. performed experiments; R.C.R. and F.S. analyzed the sequence data. D.P., R.B.D.R. and N.K. discussed results. D.P., R.B.D.R. and R.C.R. wrote the manuscript with input and commentaries from all authors.

## DATA AVAILABILITY

The raw FASTQ files from deep sequencing that support the findings of this study will be deposited (following peer-review and publication) in the Sequence Read Archive (SRA) with the primary accession code(s) <code(s) (https://www.ncbi.nlm.nih.gov/sra)>. Additional data that support the findings of this study are available from the corresponding author upon reasonable request. The raw and processed sc-RNAseq data generated in this study will be deposited in the Gene Expression Omnibus under accession number ---.

