## Supplementary material for "A method for polyclonal antigen-specific T cell-targeted genome editing (TarGET) for adoptive cell therapy applications": TarGET_Supplemental

SUPPLEMENTAL INFORMATION

**Supplemental Figures**

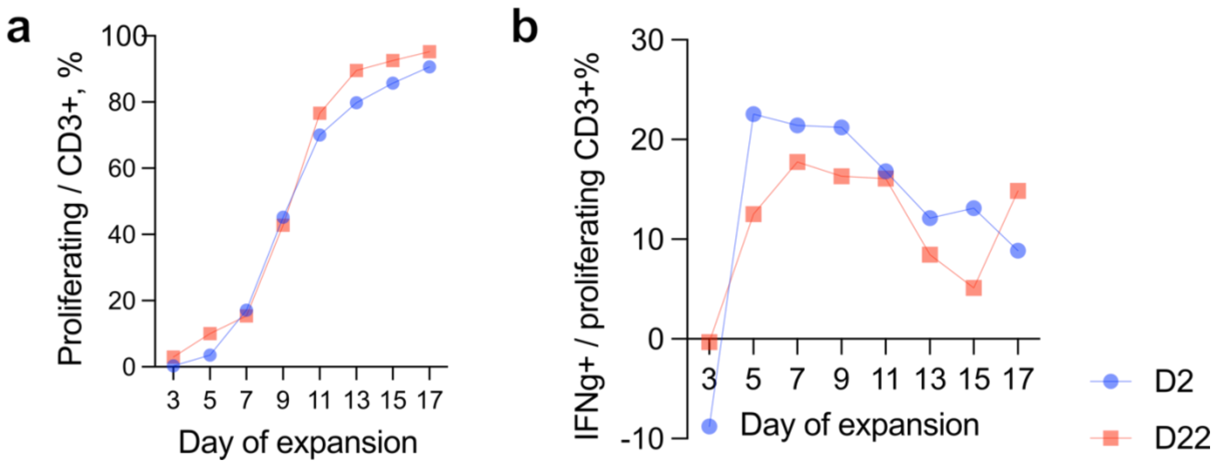


**Supplemental Fig.1: EBV-specific T cell expansion detected by cell proliferation assay. a**, dynamic of overall proliferation of CD3+ cells (proliferating cells = cells proliferated at least once). **b**, dynamic of EBV-specific T cells identified by immunocytochemistry following EBV pepmix re-stimulations.

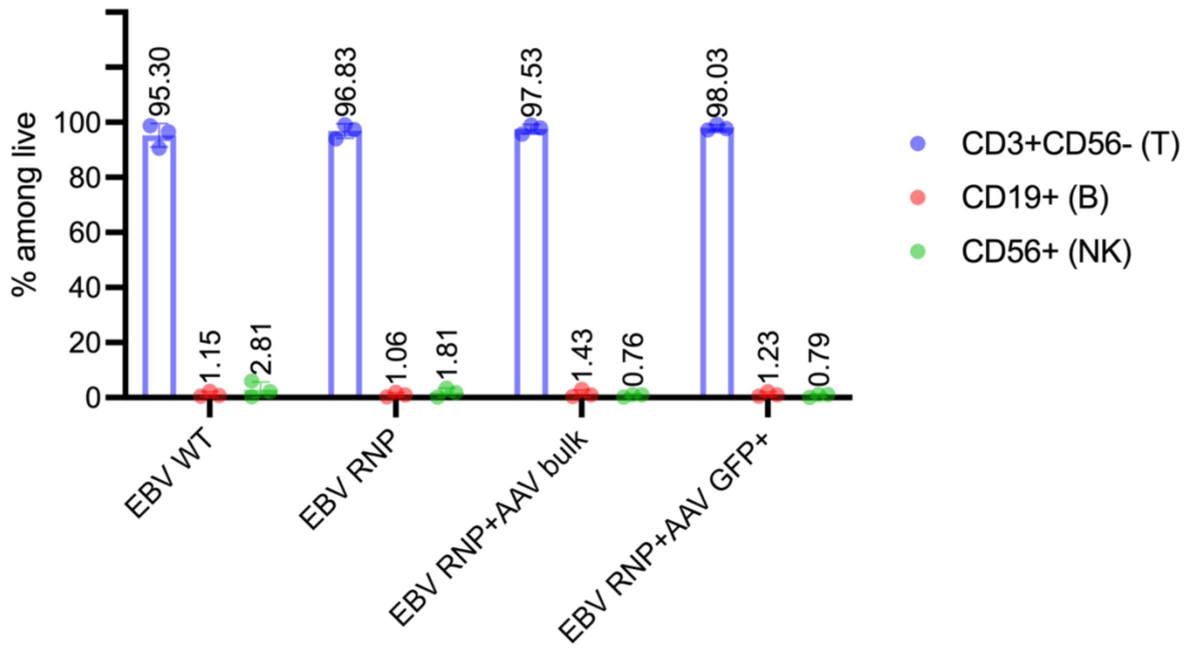


**Supplemental Fig. 2: The purity of expanded EBV-CTLs.**

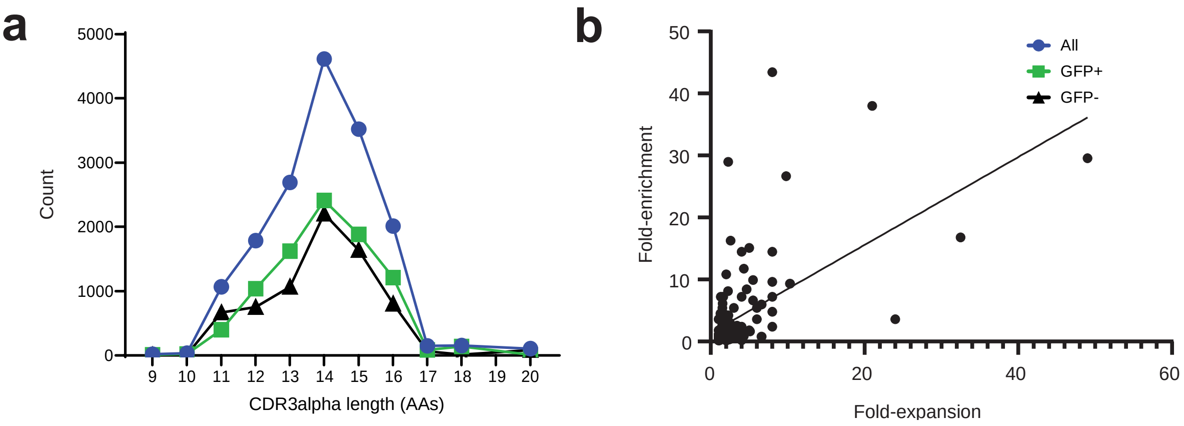


**Supplemental Fig. 3: a**, Distributions of the CDR3 alpha lengths for GFP+ and GFP- samples. Like the distributions of the CDR3 beta lengths, they do not differentiate between samples. **b,** Scatter plot of the fold-expansion and fold-enrichment values for all GFP+ clonotypes with barcodes. A linear regression of the data reveals a statistically significant correlation ($P < 0.0001$, $R^2 = 0.34$), indicating that a greater fold-expansion of the T cell clone is linked to a greater fold-enrichment in the GFP+ subset.

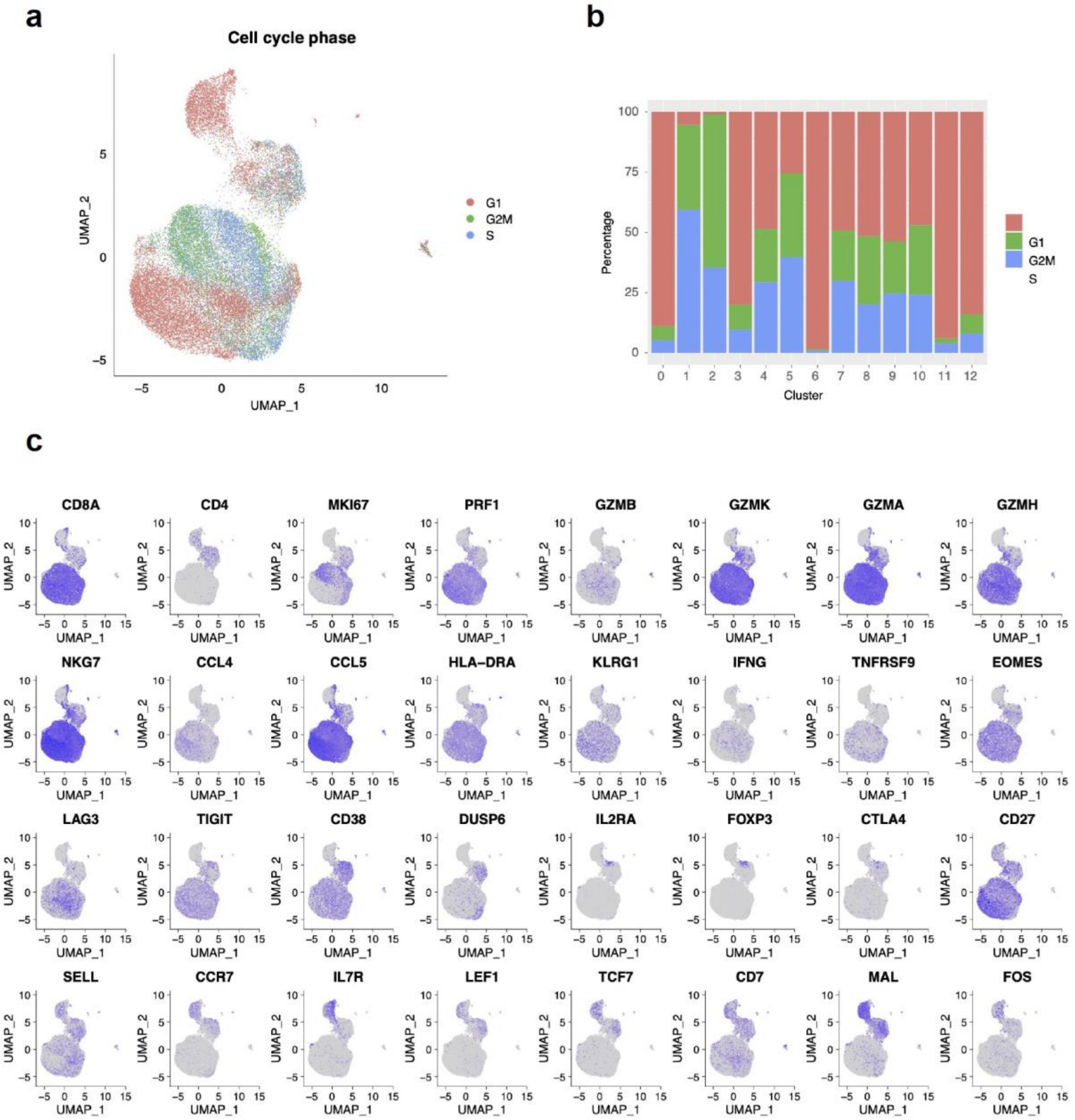


**Supplemental Fig. 4: Gene expression profiles of EBV pulsed T cells. a,** UMAP plot depicting cell cycle phase prediction for each single cell and **b,** abundance of each cell cycle phase within different clusters. **c,** projection of differential marker expression.

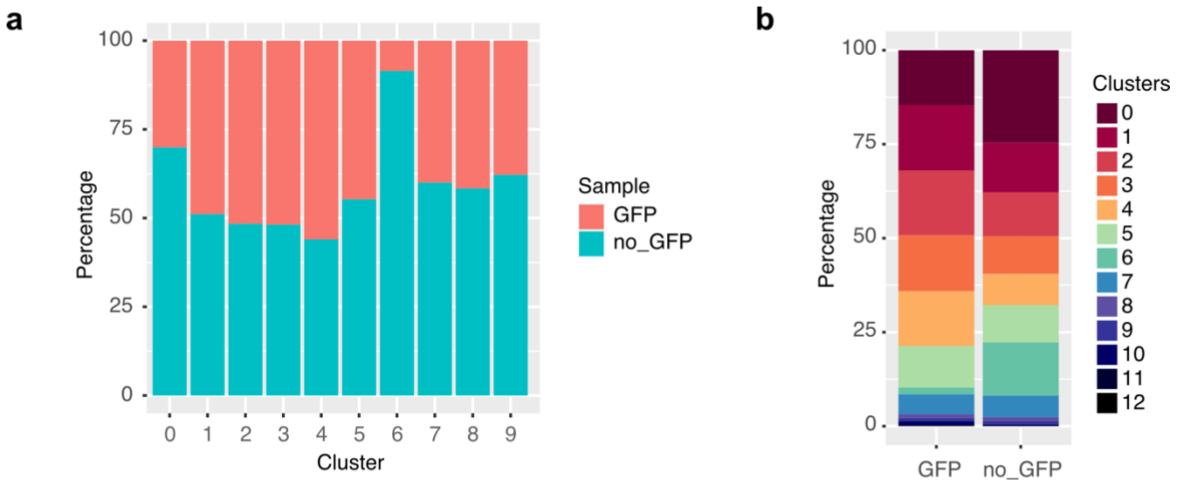


**Supplemental Fig. 5. Cluster distribution within samples. a,** Relative distribution of each cluster between GFP and no_GFP samples. **b,** distribution of clusters within each sample.

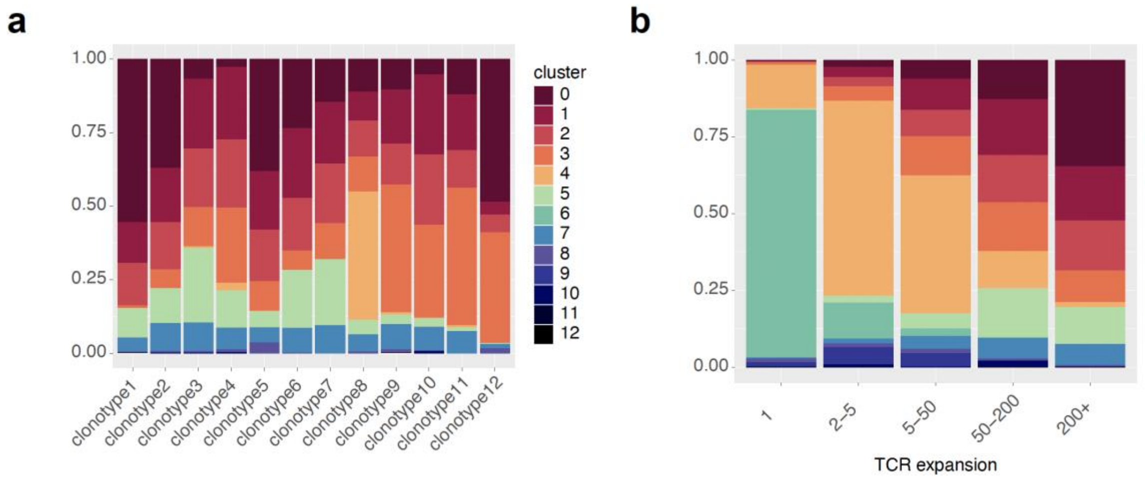


**Supplemental Fig. 6. Cluster distribution across clonotypes with different expansion rates. a,** Relative distribution of each cluster between the top 12 most expanded clonotypes. **b,** Distribution of clusters within clonotypes belonging to 5 different expansion bins.

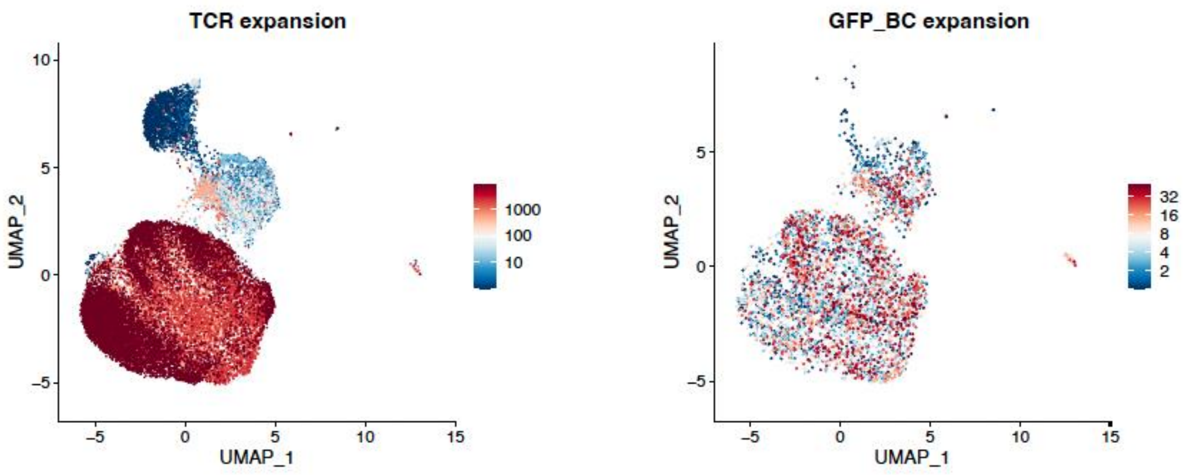

**Supplemental Fig. 7. Distribution of TCR and GFP-barcode clonotypes according to the level of expansion.**

### Supplementary Tables

**Supplemental Table 1:**

| | | |
|---|---|---|
| RB202 | GFP_end_R | TTACTTGTACAGCTCGTCCATGCCG |
| RB203 | BAR_MCS_F | NNNNNNNNNNNNGAATTCGATATCAAGCTTGTCGACC |

| | | |
|---|---|---|
| RB198 | CCR5_LHA_start_F | TTCTTTGTGGGCTAACTCTAGCGTC |
| RB199 | CCR5_RHA_end_R | GGCCAAAGAATTCCTGGAAGGTGTT |

| | | |
|---|---|---|
| RB214 | EGFP-C_Illu | CCCTCCTTTAATTCCCCATGGTCCTGCTGGAGTTCGTG |
| RB215 | EBF-rev_Illu | GAGGAGAGAGAGAGAGGTGGTTTGTCCAAACTCATC |
